# Putative NAD(P)-binding Rossmann fold protein is involved in chitosan-induced peroxidase activity and lipoxygenase expression in Physcomitrella

**DOI:** 10.1101/2023.04.20.537753

**Authors:** Eeva M. Marttinen, Eva L. Decker, Petra Heinonen, Ralf Reski, Jari P.T. Valkonen

## Abstract

Oxidative burst, the rapid production of high levels of reactive oxygen species (ROS) in response to external stimuli, is an early defense reaction against pathogens. The fungal elicitor chitosan causes an oxidative burst in the moss Physcomitrella (new species name: *Physcomitrium patens*) mainly due to the peroxidase enzyme Prx34. To better understand the chitosan responses in Physcomitrella, we conducted a screen of part of a *P. patens* mutant collection to isolate plants with less peroxidase activity than wild-type plants after chitosan treatment. We isolated a *P. patens* mutant that affected the gene encoding NAD(P)-binding Rossmann fold protein (hereafter, Rossmann fold protein). Three Rossmann fold protein-knockout (KO) plants (named Rossmann fold KO-lines) were generated and used to assess extracellular peroxidase activity and expression of defense-responsive genes including alternative oxidase (AOX), lipoxygenase (LOX), NADPH-oxidase (NOX) and peroxidase (Prx34) in response to chitosan treatment. Extracellular (apoplastic) peroxidase activity was significantly lower in Rossmann fold KO-lines than in wild-type plants after chitosan treatments. Expression of the LOX gene in Rossmann-fold KO-plants was significantly lower before and after chitosan treatment when compared to WT. Peroxidase activity assays together with gene expression analyses suggest that the Rossmann-fold protein might be an important component of the signaling pathway leading to oxidative burst and basal expression of the LOX gene in Physcomitrella.

## Introduction

Plants have developed both early and late defense reactions against pathogens (Wojtaszek 1997). Plant plasma membrane-localized pattern-recognition receptors (PRRs) perceive pathogen-associated molecular patterns (PAMP) including fungal chitin and bacterial flagellar and peptidyl glucans inducing early defense responses during PAMP-triggered immunity (PTI) (Jones & Dangl 2006; Couto & Zipfel 2016; Kawasaki et al. 2017). This very early response, which occurs within 1−5 min after PRR activation, is based on mainly the activation of pre-existing components in the plant cells (Wojtaszek 1997) and includes alkalization of the growth medium due to changes in ion fluxes across the plasma membrane (Boller 1995; Nürnberger et al. 2004; Wendehenne et al. 2002), activation of mitogen-activated protein kinase (MAPK) cascades (Asai et al. 2002), changes in protein phosphorylation (Nühse et al. 2007, Benschop et al. 2007) and oxidative burst (Apel & Hirt 2004, Torres 2010). The later defense response during PTI, which occurs within 5-30 min after PRR activation, includes the activation of defense-related genes (Boller & Felix 2009).

Early defense reactions such as oxidative burst and activation of MAPK cascades also occur among mosses. In Physcomitrella, fungal and bacterial PAMPs activate MAPK cascades that are important for the expression of defense-related genes and accumulation of cell wall−associated depositions (Bressendorff et al. 2016). Chitosan, a fungal cell wall derivate, induces peroxidase activity in *P. patens* and an immediate burst of ROS in the culture media of *P. patens* and the peat moss *Sphagnum capillifolium* (Lehtonen et al. 2009). This oxidative burst is dependent on redox homeostasis (Lehtonen et al. 2012) and involves the mitochondrial protein TSPO (Frank et al. 2007). Increase in extracellular peroxidase activity after chitosan treatment and the oxidative burst are highly dependent on peroxidase Prx34 in Physcomitrella, as *P. patens* Prx34 knockout lines do not show peroxidase activity or an oxidative response elicited by chitosan. Moreover, these mutant plants are more susceptible to fungal infection with *Fusarium* and *Irpex* spp., indicating that Prx34 is needed in defense against fungi (Lehtonen et al. 2009, Lehtonen et al. 2012).

In addition to early defense responses, the induction of defense- and ROS-related genes occurs in mosses as a response to pathogens or pathogen elicitors. Ponce de León et al. (2007) showed that inoculation of *P. patens* with culture filtrates from *Erwinia carotovora* (new species name: *Pectobacterium carotovorum*) and *Botrytis cinerea* induces the expression of the defense-related genes pathogen related-1 (PR-1), phenylalanine ammonia-lyase (PAL), chalcone synthetase (CHS) and lipoxygenase (LOX). Lehtonen et al. (2012) showed that chitosan treatment induces the expression of alternative oxidase (AOX), NADPH-oxidase (NOX) and LOX in *P. patens*. Although these studies have shown that the induction of defense- and ROS-related genes occurs in mosses as a response to pathogen or pathogen elicitors, little is currently known about signaling cascades leading to the activation of early defense responses in mosses.

Mutant collections are valuable tools for gene function studies and functional genomics. A mutant collection of Physcomitrella plants was established in the early 2000s (Egener et al. 2002, Schween et al. 2005). There were two different approaches for cDNA-based gene disruption library transformation. Egener et al. (2002) described a batch transformation in which normalized cDNA from an amplified protonema library was mutagenized and used to transform Physcomitrella. Schween at al. (2005) illustrated another method in which defined batches of 20 different mutagenized cDNAs were pooled and protoplasts were transformed with these cDNA mixtures (c20 minibatch). The collection of Physcomitrella mutants consists of more than 73,000 plants and is a valuable resource for mutant screens and identification of gene functions (Schween et al. 2005).

In this study, a part of the Physcomitrella mutant collection was screened to elucidate the pathway of peroxidase activity in response to chitosan treatment. We presumed that transgenic plants that do not respond to chitosan treatment by increased apoplastic peroxidase activity would lack a component that is involved in rapid ROS production by peroxidase in *P. patens*. Our results show that a Rossmann fold protein is involved as a novel player in the pathway leading to increased peroxidase activity after chitosan treatment and normal expression of LOX.

## Results

### Screening of the Physcomitrella mutant collection and plant regeneration

Altogether, 2622 individual hits from KO plants and 8209 hits from HoMi (high throughput minibatch, named as HoMi according to Hochdurchsatz Minibatch) plants were obtained from the moss database when the following requirements were used: phenotypic similarity with wild type, a similar number of gametophores as in WT plants and detectability with the *npt*II selection cassette. Hits among the KO plants were divided into 71 groups (consisting of different constructs) and hits among HoMi plants were divided into 511 pools (all with the transformation DNA construct).

Initially, one plant from each KO group and HoMi pool was retrieved from cryo-preservation over liquid nitrogen (Schulte and Reski 2004). To revive the plants, they were thawed and transferred to new medium according to the protocol from Schulte and Reski (2004). Any of the thawed plants that did not regenerate within 5 weeks (wk) were replaced by another plant from the same pool or group. In total, 713 plants were thawed, including three WT plants, 79 KO plants and 631 HoMi plants. Most of the KO plants (85.4%) regenerated within 4 wk from thawing and an additional 4.9% regenerated within 5 wk from thawing (Fig. 1a). Altogether, 8.5% of thawed KO plants did not regenerate within 8 wk. HoMi plants regenerate less as compared with KO plants: only 42.2% of HoMi plants regenerated within 4 wk, with an additional 9.2% regenerating within 5 wk from thawing (Fig. 1a). A substantial portion of HoMi plants (45.5%) did not regenerate within 8 wk.

**Figure 1.**
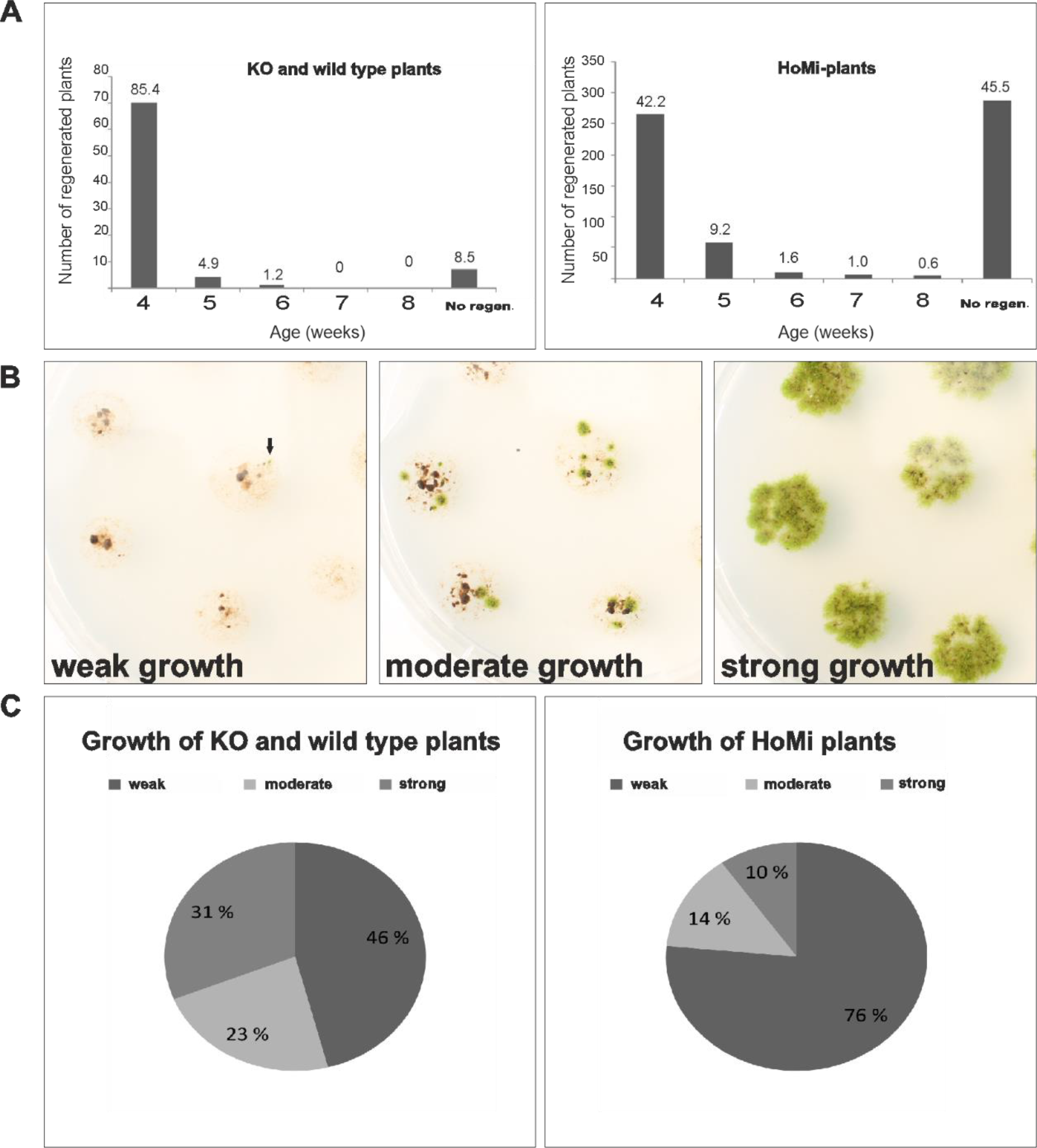
Number of regenerated transgenic Physcomitrella plants and their growth intensity. (A) Percentages of regenerated wild-type, KO (stored in liquid nitrogen) and HoMi (stored - 80° C) plants. (B) Weak (left), moderate (middle) and strong (right) growth was observed among regenerating plants. Pictures were taken eight weeks after thawing. (C) Percentages of weak, moderate and strong growth among wild-type, KO and HoMi plants.

Growth of the thawed plants varied from weak to strong (Fig. 1b). Among the thawed plants, 46% of KO plants and 76% of HoMi plants grew weakly, as indicated by only a few green dots on the agar plates. Almost one-third (31%) of KO plants and 10% of HoMi plants were growing vigorously at 4 wk after thawing (Fig. 1c).

### Analysis of transgenic Physcomitrella plants for induction of peroxidase activity after chitosan treatment

The aim of this screen was to find transgenic plants that do not show extracellular peroxidase activity after chitosan treatment. Some of the regenerated plants (34) were excluded because of their slow growth. Altogether, 385 plants were screened for their peroxidase activity upon chitosan treatment. After the first peroxidase activity assay, 149 transgenic lines that did not show peroxidase activity were chosen for a second round of screening. In the second peroxidase activity assay, 23 of those 149 transgenic plants (i.e., 6% of the initial plants) again did not show peroxidase activity upon chitosan treatment and these plants were chosen for further study.

To verify the results from the peroxidase activity screen, a third round of screening was carried out with WT plants and the 23 transgenic moss lines. The experiment was conducted twice with three independent replicates in each experiment. According to the Mann-Whitney U-test, mutant lines 57282, 58174, 54467 and 49222 had significantly lower peroxidase activity than WT plants (*P* < 0.02; Fig. 2). In all, 1.04% of the screened lines had significantly lower peroxidase activity in comparison to WT in response to chitosan treatment.

**Figure 2.**
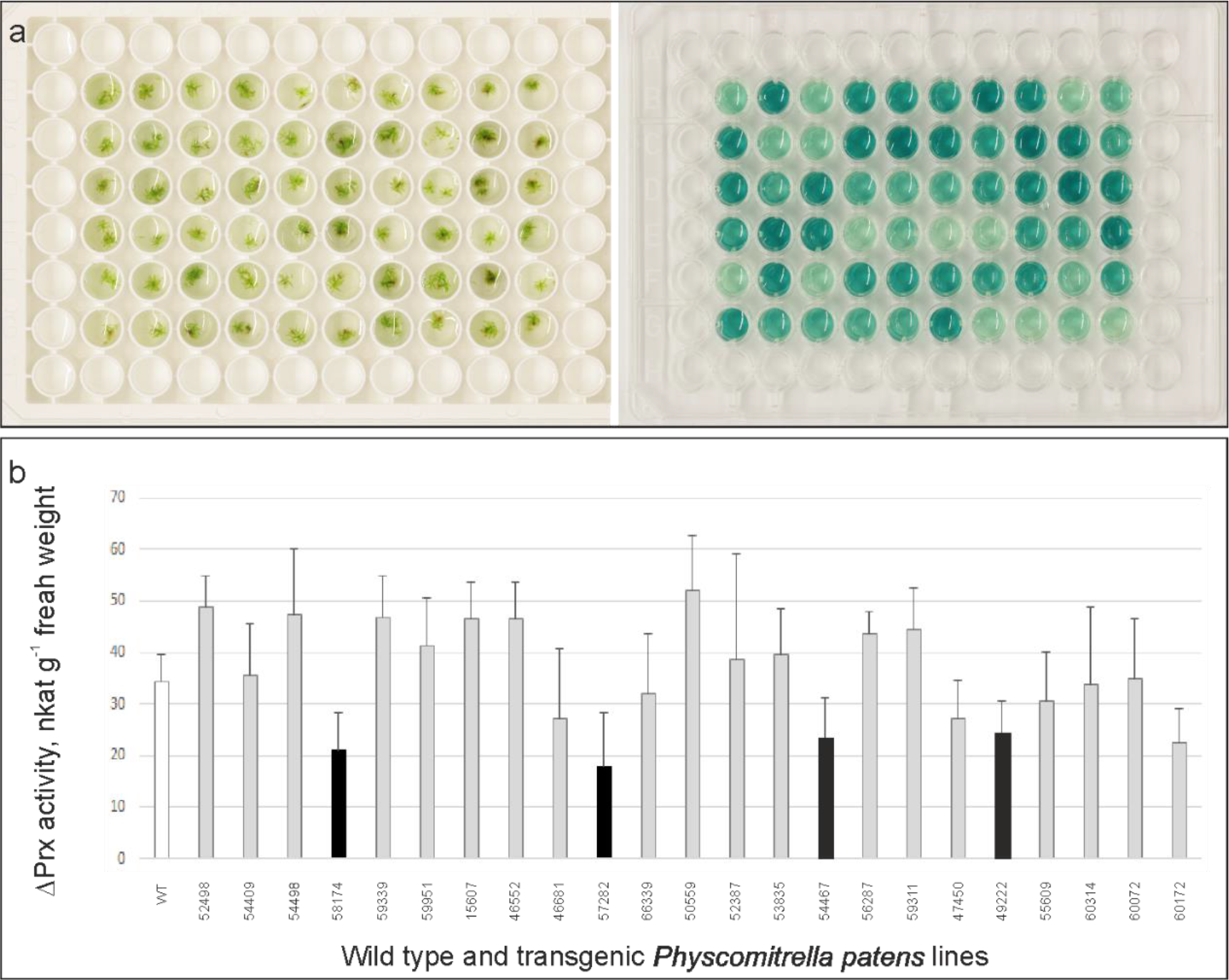
Screening of Physcomitrella mutant collection and verification of peroxidase activity. (A) An overview of Physcomitrella mutant screen on 96 well plates. (B) Peroxidase activity in wild-type and 23 transgenic *P. patens* plants after chitosan treatment. Peroxidase activity was measured after chitosan treatment in liquid culture medium. The graph shows the mean peroxidase activity from two independent experiments with three biological replicates of each transgenic moss line. Black bars indicate moss lines (57282, 58174, 54467 and 49222) that had significantly lower peroxidase activity (P < 0.02). Error bars indicate SD.

### Characterization of *npt*II integration sites

The *P. patens* line 58174 was chosen for further analyses because peroxidase activity was significantly lower than in WT in response to chitosan treatment and the activity was stable in all experiments. To investigate which genes had been mutated within the transgenic moss lines, *P. patens* line 58174 was subjected to a genome walking analysis, with WT *P. patens* as the control. The genome walking procedure resulted in the amplification of PCR products of many different sizes (*c.* 300−5000 base pairs). Sequenced PCR products were named based on the relevant restriction enzyme digest (1, *Dra*I; 2, *Eco*RV; 3, *Pvu*II; 4, *Stu*I) and the number of successfully sequenced PCR products. Consistent with the Southern blot analysis (Fig. S1), the genome walking profile of *P. patens* line 58174 indicated many *npt*II integrations (Fig. 3).

**Figure 3.**
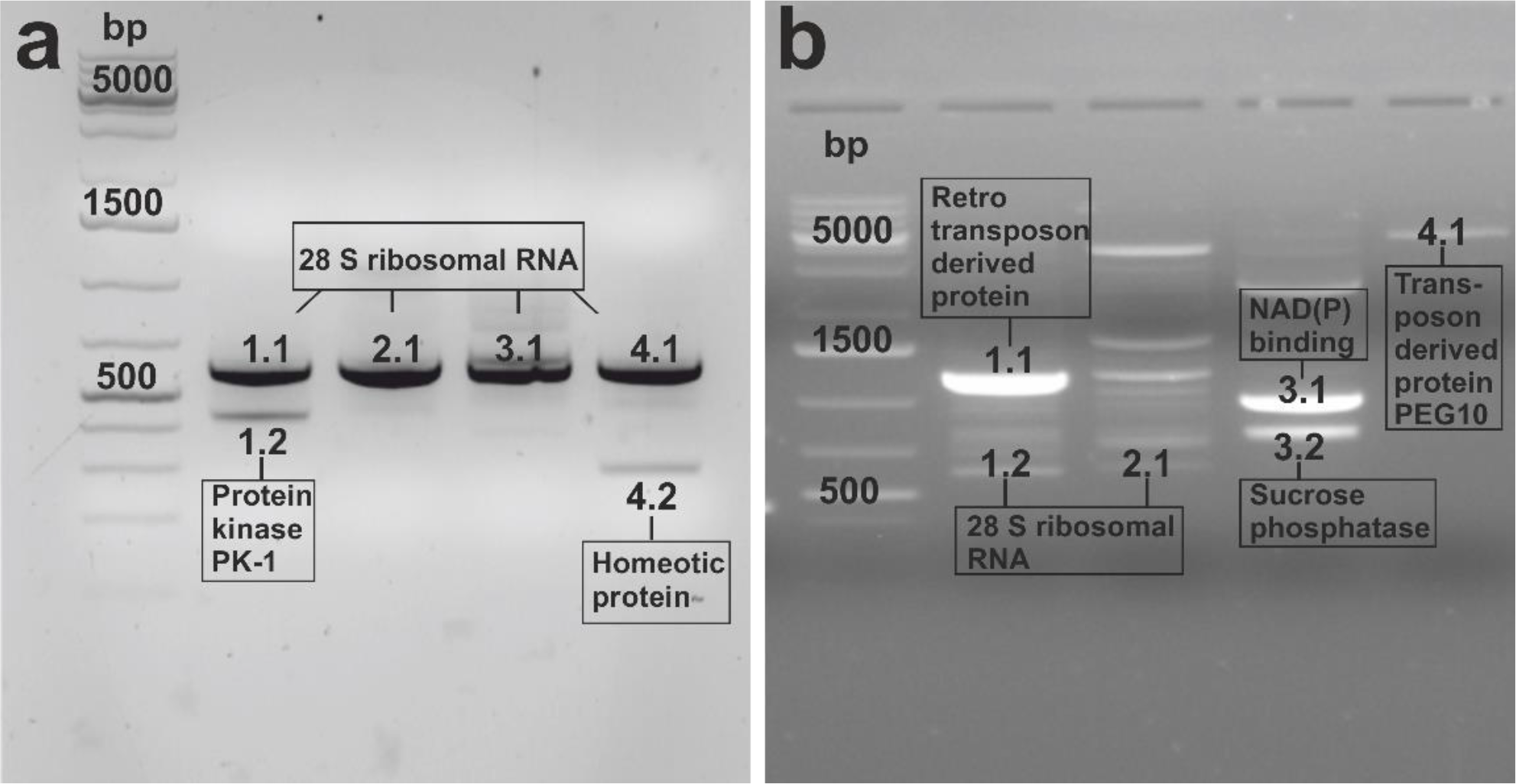
Agarose gel electrophoresis of PCR products obtained from genome walking analysis for wild-type *P. patens* (A) and the transgenic *P. patens* line 58174 (B). Sequenced PCR products were named as described in the Results. BLAST searches for nucleotide sequences were performed with the NCBI, Cosmoss and JGI *Physcomitrium patens* ssp. patens V3.3 databases. Best hits of homologs are indicated by the boxed text. A DNA ladder is shown in the first lane of each gel; the remaining lanes 1-4 correspond to DNA digested with *Dra*I, *Eco*RV, *Pvu*II and *Stu*I, respectively.

Genome walking analysis of WT *P. patens* resulted in the amplification of six clear PCR products (Fig. 3a). BLAST searches for nucleotide sequences were performed with the NCBI, Cosmoss and JGI *Physcomitrium patens* ssp. *patens* V3.3 databases. Sequencing of those products revealed that 1.1, 2.1, 3.1 and 4.1 were identical and are located in chromosome 26 at position 11844−12431. A BLAST search with the NCBI database showed that these sequences correspond to *P. patens* 28S ribosomal RNA (LOC112278192). PCR amplification product 1.2 was located in chromosome 23 at position 13952180−13952613. A BLAST search using JGI *Physcomitrium patens* ssp. *patens* v3.3 revealed that sequence 1.2 is similar to *Arabidopsis thaliana* ATP binding protein kinase gi 15218576 and to *Zea mays* protein kinase PK1 gi479356. PCR product 4.2 located in chromosome 10 at position 16420467−16420735 was predicted to be *P. patens* homeotic protein XM024530104.1.

Genome walking analysis of *P. patens* line 58174 resulted in many PCR amplification products and six of these were cloned and sequenced. PCR product 1.1 was identical to *P. patens* gene model XM00175268.5 and shared 37% identity with a retrotransposon-derived protein from the fungus *Smittium culicis* (Tuzet & Manier ex Kobayasi). PCR products 1.2 and 2.1 were identical to each other as well to PCR products 1.1, 2.1, 3.1 and 4.1 obtained from WT, corresponding to the *P. patens* 28S ribosomal RNA sequence. PCR product 3.1 was located in chromosome 26 at position 5854109−5854927. A BLAST search indicated sequence similarity with a predicted protein in *P. patens* gene model XM024511110.1. A BLASTX search revealed that the sequence was similar (62.90%) to the NAD(P)-binding Rossmann fold superfamily protein in *A. thaliana* NP 193616.2. PCR product 3.3 (766 bp) was located in chromosome 24 at position 917842−918606. A BLAST search using JGI *P. patens* ssp. *patens* v3.3 (Pp3c24_1340v3.1) revealed that this sequence was similar to *A. thaliana* sucrose phosphatase gi 30694025. PCR product 4.1 was located in chromosome 10 at position 6610509−6611403. BLASTX using the NCBI database showed a 40% sequence identity to retrotransposon-derived protein XP017945114.1 PEG10 in *Xenopus tropicalis* (Gray).

### Analysis of conserved motifs in the Rossmann fold protein homolog

The Rossmann fold protein homolog was selected for further analyses because the functional role of Rossmann fold superfamily proteins are versatile (Medvedev et al. 2021). Rossmann fold is found among nucleotide-binding proteins that bind diphosphate-containing cofactors such as NAD(H). Motif finder revealed that the protein contains 13 motifs including Rossmann fold and NmrA-like family motifs (Fig. 4a). The amino acid sequence of the *P. patens* putative (NADP)-binding Rossmann fold protein was subjected to sequence alignment and phylogenetic analyses in comparison with 15 putative NADP-binding protein homologs. The conserved region [VILWF]-X-[VIL]-X-G-X2-G-X2-[GA]-X6-[LIAFV], which is typical for Rossmann fold class III proteins (Hua et al. 2014), was detected from aligned sequences (Fig. 4b). This conserved sequence was located in the N-terminal region of the proteins, corresponding to amino acids 118−130 for *P. patens* protein model XP024366878.1.

**Figure 4.**
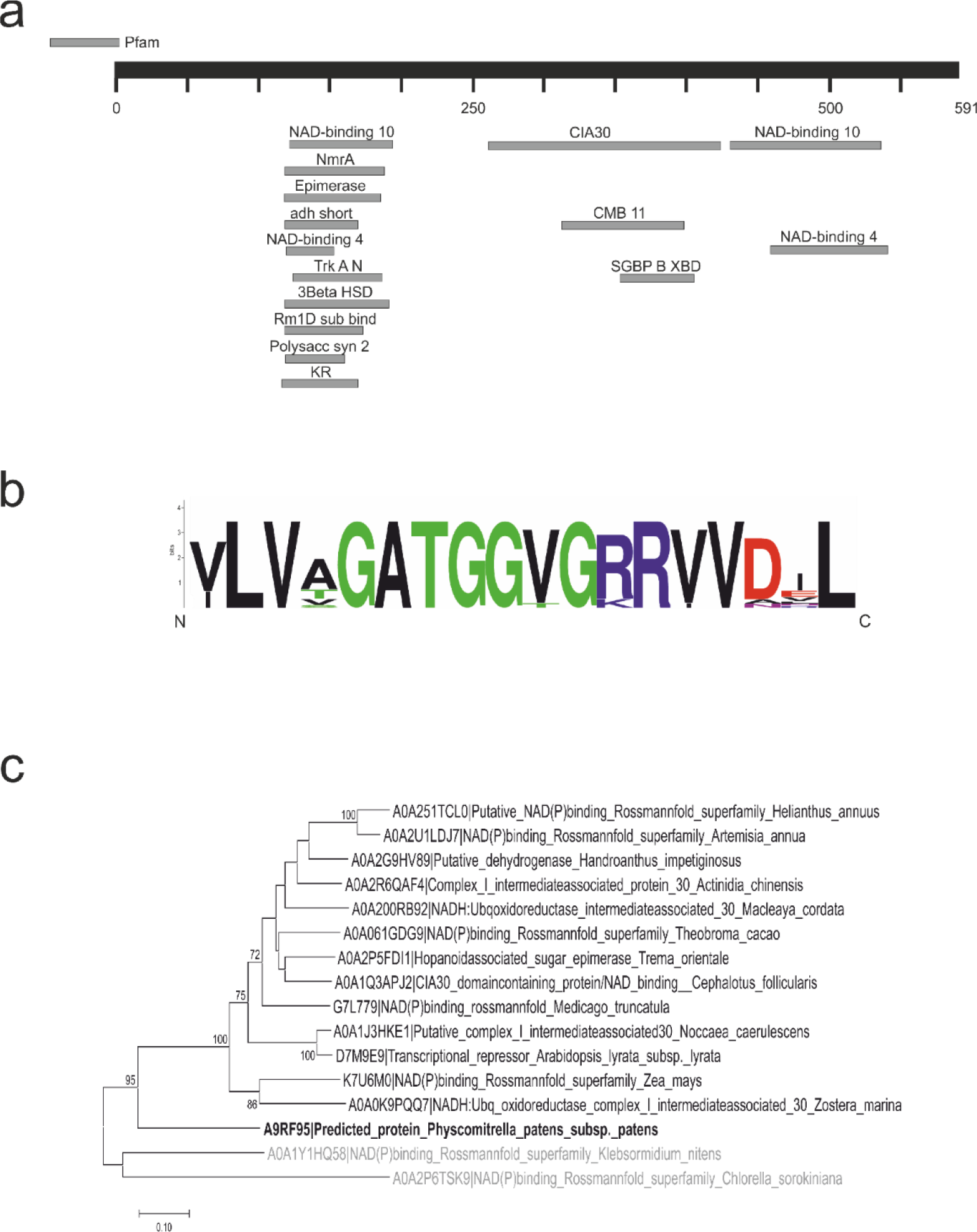
Predicted motifs and conserved region of Rossmann fold protein in *Physcomitrium patens* in Pfam database and a phylogenetic tree of selected putative NAD(P)-binding Rossmann fold proteins across other species. (A) In total, 13 motifs were found in *P. patens* XP024366878.1. NAD-binding 10 and NAD-binding 4 represent the conserved motif described below. (B) The conserved region [VILWF]-X-[VIL]-X-G-X2-G-X2-[GA]-X6-[LIAFV] typical for Rossmann fold class III proteins as visualized with the Web logo program. (C) Phylogenetic tree of putative NAD(P)-binding proteins. Phylogenetic tree based on comparison of the amino acid sequences of 16 species including two algae (gray text), one moss (bold black text) and 13 vascular plants (black text).

Aligned sequences were subjected to phylogenetic analyses. There were two main clusters observed in the maximum likelihood phylogenetic tree, algae and embryophyta, the land plants (Fig. 4c). The embryophyta cluster was divided into two main sub-clusters: the *P. patens* putative NADP-binding protein and putative NADP-binding proteins of vascular plants. In our sequence search, *P. patens* putative NADP-binding protein was the only representative from bryophytes and clustered separately from all other embryophytic NADP-binding protein homologs (Fig. 4c).

### Rossmann fold protein KO-lines have lower peroxidase activity than wild type Physcomitrella

Physcomitrella transgenic lines in which the NAD(P)-binding Rossmann fold superfamily protein was knocked out (PpRossmann fold KO-lines) were generated (Fig. S2a), taking advantage of the highly efficient gene targeting in this species (Wiedemann et al. 2018). Sequencing for the PpRossmann fold gene replacement cassette in pGEM-T vector that had been used to generate the lines was used to confirm that the correct fragments were present. Three stable gene replacement lines were chosen for further analysis. Integration of the gene replacement cassette was verified by PCR (Fig. S2b). The phenotype of the PpRossmann fold KO-lines resembled that of WT plants (Fig. S2c) and flow cytometry according to Heck et al. (2021) indicated that all lines were haploid like WT *P. patens.* Because gene targeting in Physcomitrella can occasionally generate off-target integration of the KO-construct elsewhere in the genome by illegitimate integration (Kamisugi et al. 2006, Wiedemann et al. 2018), we used three independent KO-mutant lines for further analysis.

Apoplastic peroxidase activity of the three PpRossmann fold KO-lines (17, 24 and 35) was measured to determine whether the Rossmann fold protein has a role in peroxidase activity in response to chitosan treatment. These three moss lines had significantly lower extracellular peroxidase activity than WT plants across three independent experiments (Fig. 5). Tukey’s honestly significant difference (HSD) test showed that PpRossmann fold KO-line 17 differed from WT at *P* < 0.05 and PpRossmann fold KO-lines 24 and 35 differed from WT at *P* < 0.01.

**Figure 5.**
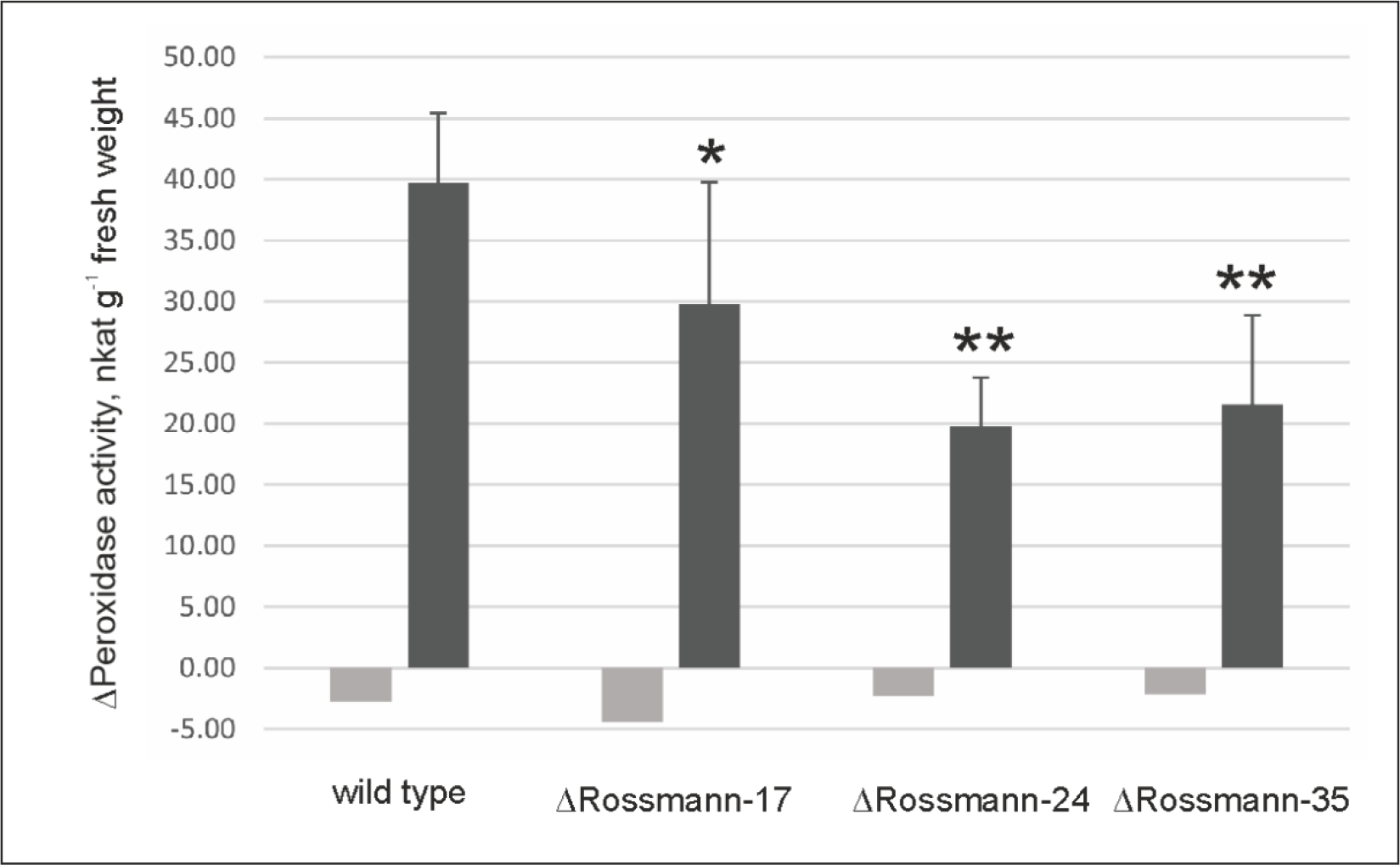
Peroxidase activity in Physcomitrella wild-type and in *P*. *patens* lines with knockout of Rossmann fold protein in response to chitosan treatment. Bars indicate the change in peroxidase activity (background peroxidase activity T10 - T0) as nanokatals per gram fresh weight measured in liquid culture medium after addition of chitosan (black) or water (light gray) in three independent experiments. Error bars represent the standard deviation of the data and asterisks indicate a significant difference relative to chitosan-treated wild type according to Tukey’s HSD test (*, P < 0.05; **, P < 0.01). ANOVA, F3,62 = 15.2.

### Lipoxygenase7 gene is constantly repressed in PpRossmann fold KO-lines

As treatment with chitosan induces the expression of the defense response genes AOX, LOX, NOX and Prx (Lehtonen et al. 2012) we analyzed the expression of these genes in PpRossmann fold KO-lines (17, 24, 35) and WT before and 40 min after chitosan treatment. In the three experiments, the transcript level of PpLOX7 was significantly repressed in all PpRossmann fold KO-lines before chitosan treatment (Fig. 6) (ANOVA F_3,12_ = 935.4, *P* < 0.001). LOX7 expression increased slightly in all PpRossmann fold lines at 40 min after chitosan treatment but was still significantly lower than in WT plants (ANOVA F_3,12_ = 9.1, *P* < 0.05 for KO-line17 and *P* < 0.01 for KO-line 24 and KO-line 35).

**Figure 6.**
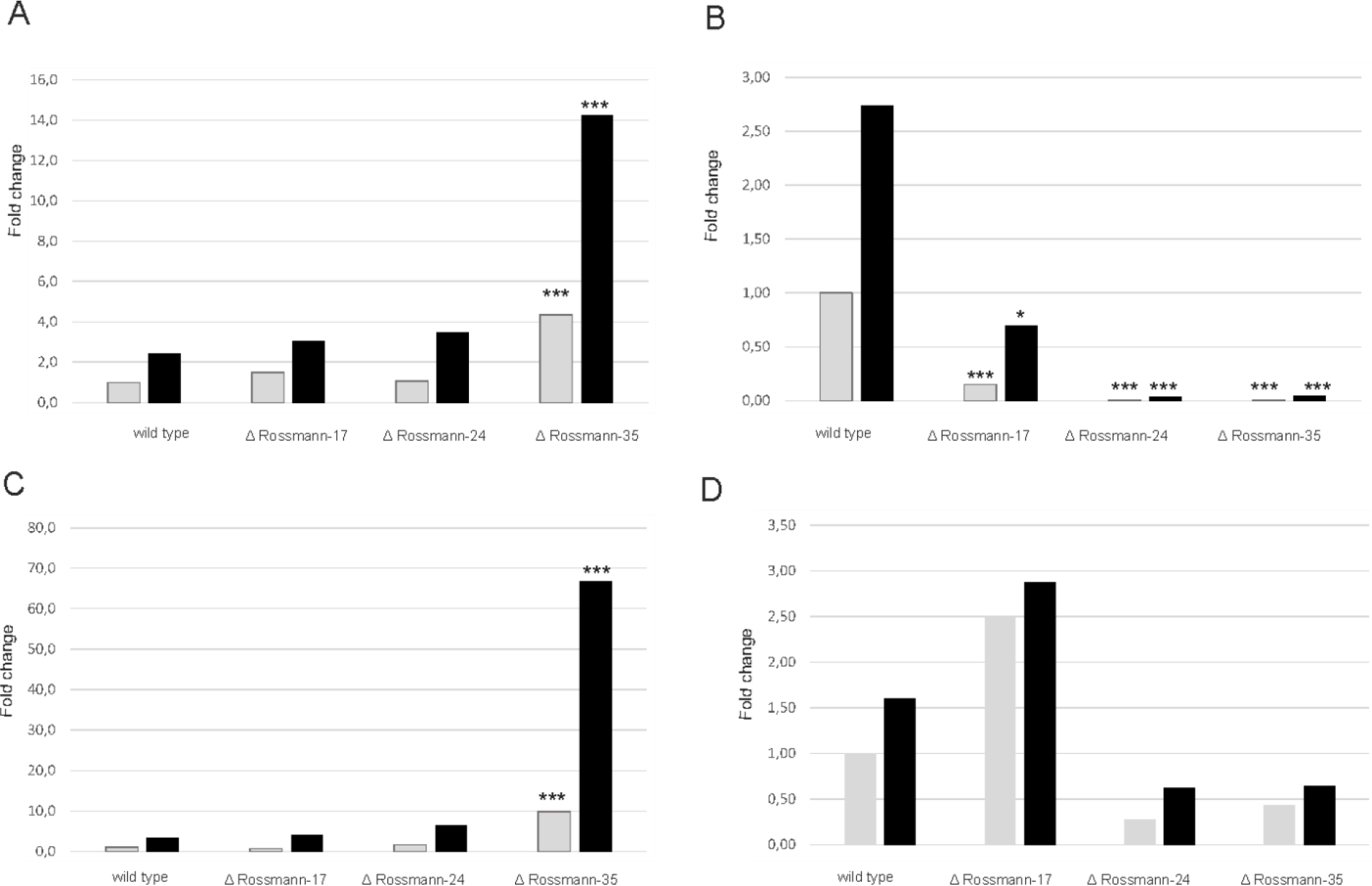
Genes as defense responsive (Leon et al. 2007) in wild-type *P. patens* as compared with expression in the Rossmann fold KO-lines (17, 24 and 35). The expression of (A) alternative oxidase, (B) lipoxygenase, (C) NADPH-oxidase, (D) peroxidase are shown before treatment (gray bars) and 40 min after chitosan treatment (black bars). Elongation factor α was used as a reference gene for normalization. The experiment was conducted three times and each experiment had two biological replicates for each treatment. qPCR experiment was conducted with three technical replicates. Asterisks indicate a significant difference relative to wild type for the specific time point (before or 40 min after treatment) according to Tukey’s HSD test (*, P < 0.05; **, P < 0.01; ***, P < 0.001). ANOVA, F3,12 = 20,459 in A, 935,424 in B, 19,446 in C and 1,070 in D at 0 min time point. ANOVA, F3,12 = 8,111 in A, 9,101 in B, 14,146 in C and 0,876 in D at 40 min time point.

In the three experiments, the transcript level of PpAOX, PpNOX and PpPrx34 increased after chitosan treatment in WT and in all PpRossmann fold KO-lines. The expression levels of AOX and NOX genes were significantly induced in one PpRossmann fold KO-line (35) before and after chitosan treatment (Fig. 6).

## Discussion

Here, we used a Physcomitrella mutant collection as a promising resource for analyzing gene function. Screening of 385 plants enabled us to find candidate plants with altered responses to chitosan treatment. Finally, 1.04% of screened plants had significantly lower extracellular peroxidase activity in comparison to WT *P. patens* upon chitosan treatment. These results indicate that the Physcomitrella mutant collection provides a promising source of novel genes for defense signaling cascades downstream of a specific elicitor.

The one plant with reduced amounts of peroxidase activity in response to chitosan that was fully analyzed here had a mutation in a gene encoding a putative Rossmann fold protein. KO-lines lacking the gene for Rossmann fold protein had significantly lower peroxidase activity upon chitosan treatment. The Rossmann fold is a common protein motif and this superfamily of proteins has versatile functions (Medvedev et al. 2021). A Rossmann fold is found in many metabolic enzymes that bind cofactors such as NAD and NADP. NADP is an essential molecule for energy metabolism and is involved in signaling pathways that regulate many cellular processes including gene transcription and apoptosis (Berger et al. 2004, Ying 2006).

Dehydrogenases can positively regulate the resistance to pathogens in vascular plants (Choi et al. 2008, Hwang et al. 2012). Choi et al. (2008) showed that the Rossman fold–containing enzyme menthone reductase1 (MNR1), a member of the short-chain dehydrogenase/reductase (SDR) superfamily, positively regulates the defense against pathogens in pepper (*Capsicum annuum*).

*CaMNR1*-silenced pepper plants were significantly more susceptible to infections by *Xanthomonas campestris* pv *vesicatoria* and *Colletotrichum coccodes*. Hwang et al. (2012) showed that SDR3 is directly involved in plant defense. Five days after spray inoculation with *Pseudomonas syringae* DC3000, the bacterial population in *Arabidopsis* plants that overexpressed SDR3 was 4-to 6-fold lower than that of wild type (Hwang et al. 2012). In addition, the bacterial population of the *Atsdr3* mutant lines was 3-to 5-fold higher than that of WT plants. Our results show that *P. patens* NAD(P)-binding Rossmann fold protein is, at least in part, required for peroxidase activation in non-vascular plants and thus it might contribute also to pathogen defense.

Further, our results show that in the absence of Rossmann fold protein the expression of LOX7 was significantly reduced, indicating that a Rossmann fold protein is essential for normal expression of LOX7 in Physcomitrella. Previous studies indicate the role of lipoxygenases in defense responses in this species. Ponce de León et al. (2007) showed that LOX expression is moderately induced in culture filtrates of *E. carotovora* −treated moss colonies at 4 and 24 h. LOX expression is also significantly increased in *B. cinerea* −inoculated *P. patens* plants. Oliver et al. (2009) showed that *Pythium debaryanum* and *Pythium irregulare* infections trigger the accumulation of PpLOX gene transcripts. LOX transcript accumulation begins at 2 h and increases up to 6−8 h after *Pythium* inoculation. LOX7 gene induction has also been observed in response to chitosan treatment (Lehtonen et al. 2012). PpLOX7 gene transcripts are slightly increased by 15 min and are significantly increased by 30 and 90 min after treatment with chitosan. Furthermore, Lehtonen et al. (2014) showed that lipoxygenase 1 is exclusively secreted in response to treatment with chitosan in *P. patens*. These results indicate that *P. patens* lipoxygenases 1 and 7 are induced by pathogens or pathogen elicitors and thus they might be important components for the basal defense of *P. patens*.

Lehtonen et al. (2009) showed that application of chitosan to suspension cultures of *P. patens* leads to a rapid increase in peroxidase activity in the medium. Furthermore, Lehtonen et al. (2014) found that Prx34 is among the most abundant proteins in the secretome of *P. patens* in control cultures and the amount increases upon chitosan treatment. In our study, the transcript level of PpPrx34 increased after chitosan treatment in WT and in all PpRossmann fold KO-lines but we found no significant differences in the transcript levels of Prx34 between WT and mutants. The difference in peroxidase activity but not in transcript level between WT and PpRossmann fold KO-lines suggests differences in the apoplastic amounts of Prx34 between WT and mutant. Thus, the Rossmann fold protein discovered here might be involved in the regulation of peroxidase secretion rather than peroxidase production upon chitosan treatment. Such a mechanism would be important for a rapid defense response of Physcomitrella. This model can be analyzed in a subsequent proteomics study.

In contrast, depletion of the Rossmann fold protein by targeted gene knockout negatively affected LOX7 gene expression at the transcript level while other defense response genes (AOX, NOX) were not affected in their transcript levels. This repression of LOX7 gene expression was detectable in untreated and in chitosan-treated mutant plants, indicating that the Rossmann fold protein discovered here is part of a signaling cascade relevant for the expression of the defense response gene LOX7.

Taken together, screening of a Physcomitrella mutant collection for reduced extracellular peroxidase activity lead to the discovery of a Rossmann fold protein in this moss. This protein is a novel factor in the signaling cascades leading to increased extracellular peroxidase activity in response to chitosan treatment and to normal expression of LOX7, while two other defense response genes tested here (AOX, NOX) remained unaffected. These results indicate a complex regulation of the defense response against pathogens in Physcomitrella.

Recently, it was shown that ROS treatment of Physcomitrella induced systemic electrical and calcium waves that transmit this information to distant parts of the plant, resulting in altered gene expression there (Koselski et al. 2023). Similar experiments can help to dissect the precise role of the Rossmann fold protein described here in the signaling cascade of this moss upon pathogen infection. Moreover, our mutant screen and genome walking has identified several more candidate proteins that might be involved in the signaling events leading to oxidative burst upon chitosan treatment in Physcomitrella. These candidate proteins can be analyzed in subsequent experiments by targeted gene knock outs as outlined here.

## Materials and methods

### Selection of plants to screen from the Physcomitrella mutant collection

Plants from the *Physcomitrella patens* (Hedw.) B.S.G. (new botanical name: *Physcomitrium patens* (Hedw.) Mitt.) mutant collection (Egener et al. 2002, Schween et al. 2005) were screened to find transgenic moss plants that differ in their response to the fungal elicitor chitosan. Moss database (moss db) (Schween et al. 2005) was used to identify plants for this screen based on three requirements: the selected transgenic plants were similar to WT plants (group A), had the same number of gametophores as WT plants and had the *npt*II selection marker integrated within their genome. Plants from two different types of transformation were used: KO and cDNA minibatch plants (high throughput minibatch, named as HoMi according to Hochdurchsatz Minibatch). HoMi plants have been made so that several (20) transposon-tagged cDNAs are co-transformed in one experiment, each of which was repeated five times (Schween et al. 2005). Only those HoMi plants whose gametophore cDNA library had been mutated were included for the search. All of the WT plants and the mutant plants were stored in liquid nitrogen and most of the HoMi plants in a – 80 °C freezer.

Selected mutant plants and three WT plants were thawed on ice and placed on solid complete medium (1.85 mM KH_2_PO_4_, 3.35 mM KCl, 1mM MgSO_4_ · 7H_2_O, 4.23 mM Ca(NO_3_)_2_ · 4 H_2_O, 0.05 mM FeSO_4_ · 7 H_2_O, 0.05 mM H_3_BO_3_, 0.05 mM^1^ MnSO_4_ · 1 H_2_O, 0.015 mM ZnSO_4_ · 7 H_2_O, 2.5 µM KI, 0.59 µM Na_2_MoO_4_ · 2 H_2_O, 0.05 µM CuSO_4_ · 5 H_2_O, 0.8 µM Co(NO_3_)_2_ · 6 H_2_O [pH adjusted to 5.8 using KOH], 22.2 µM myoinositol, 20.1 µM choline chloride, 8.12 µM nicotinic acid, 1.5 µM thiamine-HCl, 1.5 µM pyridoxine, 0.04 µM biotin, 1.82 µM *p*-aminobenzoic acid, 3.99 µM Ca-D-pantothenate, 0.04 µM l^−1^ riboflavine, 50 µM adenine, 13.8 µM Na-palmitinic acid, peptone (250 mg l^−1)^, 5 mM ammoniumtartrate and 278 mM glucose) (Egener et al. 2002). Moss plants were cultivated in a growth cabinet at 23 °C under a 16 h: 8 h, light: dark photoperiod. Regeneration of plants was recorded after 4 wk from thawing and continued weekly until plants were 8 wk old. Growth intensity was monitored during the regeneration. After regeneration, plants were transferred to solid Knop medium supplemented with microelements (Reski & Abel 1985, Egener et al. 2002) and cultivated in the growth chamber as described above.

### Screen of transgenic plants: chitosan treatment and peroxidase activity assay

After regeneration, WT and transgenic plants were grown on Knop medium supplemented with microelements until gametophores were apparent. A few gametophores, typically two to four, were transferred to 96-well plates containing 350 µl of BCD medium (1 mM MgSO_4_, 1.85 mM KH_2_PO_4_ (pH 6.5 adjusted with KOH) 10mM KNO_3_, 45 µM FeSO_4_, 0.22 µM CuSO_4_, 0.19 µM ZnSO_4_, 10 µM H_3_BO_4_, 0.10 µM Na_2_MoO_4_, 2 µM MnCl_2_, 0.23 µM CoCl_2_, 0.17 µM KI (Ashton and Cove, 1977) supplemented with 1 mM CaCl_2_, 45 µM Na_2_EDTA and 5mM ammonium tartrate ((NH_4_)_2_C_4_H_4_O_6_). Moss plants were cultivated in the growth cabinet at 23 °C under a 16 h: 8 h, light: dark photoperiod. Regeneration of plants was recorded after 4 wk from thawing and continued weekly until plants were 8 wk old. Growth intensity (weak, moderate or strong growth) was monitored during the regeneration. After regeneration, plants were transferred to solid Knop medium supplemented with microelements and cultivated in the growth chamber as described above.

Few gametophores, typically two to four were transferred to 96 well plates containing 350 µl of BCD broth (Ashton & Cove 1977). To minimize the edge effect, the outermost wells were filled with BCD broth only. Plants were grown in 96-well plates under a transparent lid for 4 d under the stationary condition as described above.

After this growth period, plants were subjected to chitosan or control treatments followed by the peroxidase activity assay. Three replicates of each plant were treated with chitosan and three replicates were treated with water. A stock solution of chitosan (10 mg ml^−1^; MW *c.* 2000−5000; Wako Pure Chemical Industries Ltd, Osaka, Japan) was prepared by dissolving chitosan in distilled water and this solution was then pre-filtered twice thought a 1.6-µm filter (Whatman, Maidstone England) and was sterile-filtered with a 0.2-µm filter (VWR, USA). Background samples of BCD broth (10 µl) were taken before chitosan or control treatments and placed on ice. Thereafter, the stock solution of chitosan was added to a final concentration of 0.5 mg ml^−1^. An equal volume of sterile water was used for the control treatment. Samples of BCD broth supplemented with water or chitosan (10 µl) were taken at 5 and 30 min after treatment and placed on ice.

The peroxidase activity assay was performed as described (Lehtonen et. al. 2009) with slight modifications. Briefly, the peroxidase reaction mixture contained 0.5 mM 2,2’-azino-bis(3-ethylbenzothiazoline-6-sulfonic acid) (ABTS) and 100 µM H_2_O_2_ in phosphate-citrate buffer (21 mM phosphate, 17 mM citrate, pH 4.0). For the peroxidase activity assay, 100 µl of reaction mixture was added to the samples in 96-well plates and mixed by pipetting. The reaction mixture was incubated for 60 min at room temperature and absorbance at 405 nm was measured by a microplate reader. Absorbance differences between background samples and treated samples were calculated. Moss lines that did not show peroxidase activity after chitosan treatment at either time point (5 min and 30 min after treatment) were screened again for confirmation.

### Verification of screening results

The peroxidase activity assay was performed once more to verify the results from screening and for generated KO-lines as described by Lehtonen et al. (2009). Briefly, protonemal tissue samples (1 wk old) from 23 transgenic moss lines and a WT line were individually homogenized in sterile water and placed on cellophane-covered BCD medium solidified with 0.8% agar. After two weeks moss cultures were transferred to BCD plates without ammonium tartrate and were collected 1 wk later from an area of 2.25 cm^2^. Tissue was transferred to 2 ml of BCD broth supplemented with 5 mM ammonium tartrate and 1% glucose in 15-ml conical tubes. Tubes were agitated in a rotary shaker (160 rpm) for 1 wk at 25 °C under a 12 h: 12 h, light: dark photoperiod with light intensity of 160 µmol m^−2^ s^−1^. Moss suspension cultures were treated with sterile water (control) or chitosan to reach a final concentration of 0.5 mg ml^−1^. Samples of BCD broth (50 µl) were collected before any treatments (T0) and 10 min after chitosan or water treatment (T10) and were placed on ice. The peroxidase activity reaction mixture contained 0.5 mM ABTS and 200 µM H_2_O_2_ in phosphate-citrate buffer (29 mM phosphate, 23 mM citrate, pH 4.0). Absorbance was measured at 405 nm using a spectrophotometer (UV-1700; Shimazu, Kyoto, Japan) at room temperature. The experiment was conducted twice with three independent replicates for each plant and each treatment. Activity was calculated based on oxidation of ABTS as katals (mol s^−1^) per gram of fresh moss tissue. The millimolar extinction coefficient of oxidized ABTS at 405 nm is 36.8.

### Statistical analyses

The Mann-Whitney U-test (IBM SPSS Amos 24) was chosen to measure if there were statistical differences with respect to peroxidase activity among the WT moss and 23 transgenic moss lines. Each mutant plant was compared with WT. A significance level of 0.02 was used. The null hypothesis was that there is no difference in peroxidase activity between a mutant plant and WT plant after chitosan treatment.

### DNA extraction

DNA was extracted from *P. patens* according to Ausubel et al. (1995). Briefly, moss protonemal tissue (*c.* 100 mg) was frozen in liquid nitrogen and ground to a powder. DNA extraction was carried out with 500 µl of cetyl trimethyl ammonium bromide (CTAB) buffer (Ausubel et al. 1995) and incubated at 65 °C for 1 h. Samples were extracted with chloroform and centrifuged at 10000 × *g* for 15 min at 4 °C. The aqueous phase was collected and DNA was precipitated with 500 µl of isopropanol. The DNA was pelleted by centrifugation (10000 × *g* for 10 min at 4 °C) and washed with 70% ethanol. The pellet was re-suspended in 500 µl of TE buffer (10 mM Tris-HCl, pH 8.0; 1 mM Na_2_-EDTA) and treated with RNaseA at 60 °C for 1 h (1 µg ml^−1^, Sigma).

### Southern blot analysis

Southern blot analysis was carried out according to the DIG Application Manual (Roche, Mannheim, Germany). In brief, genomic DNA (10 µg) was digested with 10 µl (100 U) *Pvu*II restriction enzyme overnight and separated on a 1% agarose gel by electrophoresis. Separated DNA fragments were transferred to MANGA nylon transfer membrane (Osmonics Inc., New York, USA) by capillary transfer overnight and cross-linked with ultraviolet light. A probe was designed to bind to the region within the neomycin phosphotransferase II (nptII) selection cassette. Forward (5’-gaggacgtactcggatggaa-3’) and reverse (5’-aatatcacgggtagccaacg-3’) primers (2.5 µl, 20 µM) were used for amplification of DIG-labeled probe. A PCR reaction (50 µl) that included 1.25 µl (6.25 U) of Taq polymerase (ThermoFisher Scientific, Vilnius, Lithuania); 5 µl 10× Taq PCR buffer; 2 µl MgCl_2_ (25 mM); 1 µl template DNA; 1 µl each of dATP, dCTP, dTTP (10 mM) and 0.65 µl of DIG-dUTP (1 mM) was used. The final reaction volume was adjusted to 50 µl with nuclease-free water. Prehybridization for 60 min and overnight hybridization were carried out with a DIG-labeled probe (1 µg) at 65 °C with DIG Easy Hyb Granules (Roche). A chemiluminescent assay was used to visualize the probe on the membrane. Alkaline phosphatase−labeled anti-digoxigenin and CDP-Star (Roche) substrate were used for detection of probe.

### Genome walking

The Universal GenomeWalker 2.0 kit (Clontech Laboratories Inc., now Takara Bio Inc., California, USA) was used to determine the sequence adjacent to *npt*II. For library generation, four genomic DNA samples (0.5 µg each) were individually digested with *Dra*I, *Eco*RV, *Pvu*II and *Stu*I restriction enzymes (80 U each) and ligated to genome walker adaptors according to the manufacturer’s instructions. Gene-specific primers were designed in the region of *npt*II. A gene-specific outer primer (GSP outer right; 5’-tcctctccaaatgaaatgaacttcc-3’) and gene-specific nested primer (GSP nested right; 5’-cttccttttctactgtcctttcgatg-3’) were used in combination with genome walker AP1 (GSP1 outer) and genome walker AP2 (GSP2 nested) primers provided with the kit.

Primary PCR contained 19.5 µl of deionized water, 2.5 µl of 10× Advantage 2 PCR buffer, 0.5 µl dNTPs (10 mM each), 0.5 µl AP1 (10 µM), 0.5 µl GSP outer primer (10 µM), 0.5 μl Advantage 2 Polymerase Mix (50×) and 1 µl of each digested DNA library (digested with *Dra*I, *Eco*RV, *Pvu*II and *Stu*I). The PCR cycle consisted of 94 °C for 25 s and 72 °C for 3 min for seven cycles, followed by 94 °C for 25 s and 67 °C for 3 min for 32 cycles and 67 °C for an additional 7 min. Secondary PCR contained 19.5 µl of deionized water, 2.5 µl of 10× Advantage 2 PCR buffer, 0.5 µl dNTPs (10 mM each), 0.5 µl AP2 (10 µM), 0.5 μl GSP nested (10 µM), 0.5 μl Advantage 2 Polymerase Mix (50×) and 1 µl of primary PCR product, diluted 1:10.

PCR products were analyzed on a 1.5% agarose gel, purified with an E.Z.N.A. Gel Extraction kit (Omega Bio-tek Inc., Georgia, USA) and cloned into the pGEM-T easy vector (Promega, Madison, WI, USA) according to instructions, except that 0.5 µl of pGEM-T easy vector (50 ng µl^−1^), 0.5 µl of T4 DNA ligase (3 Weiss U µl^−1^) and 1.5 µl of purified PCR products were used. Ligation mixture was incubated at 4 °C overnight and 2.5 µl was used for transformation of *Escherichia coli* DH5α competent cells. Plasmids were extracted with the Gene Elute Plasmid Miniprep kit (Sigma, St. Louis, USA) and sent for sequencing to two different companies (EuroFins Genomics, Germany, Macrogen The Netherlands).

### Motif search and phylogenetic analyses

*P. patens* amino acid sequence Pp3c26_9450V3.1 was subjected to motif finder to investigate putative motifs (GenomeNet, Kyoto University Bioinformatics Center). Default settings were used for the motif search.

The UniProt databaseKB was searched for homologous proteins of the putative *P. patens* NAD(P)-binding Rossmann fold superfamily protein (*P. patens* gene model XM_024511110.1, XP_024366878.1) with default settings. Sequences that scored >1.7 and had a putative annotation were chosen for alignment. Alignment was performed with ClustalW (Thompson et al. 1994) with default parameters: Gonnet protein weight matrix, Gap opening penalty = 10, Gap extension penalty for pairwise alignment = 0.1 and for multiple alignment = 0.2. Conserved regions were visualized with the WebLogo program (Crooks et al. 2004). Phylogenetic analyses using the maximum likelihood method with 1000 bootstrap replicates and a WAG model were performed using MEGA version 7.0.14 (Kumar et al. 2016).

### Knockout constructs

The gene encoding NAD(P)-binding Rossmann fold superfamily protein Pp3c26_9450V3.1 was used to construct knockout mutant lines according to Strepp et al. (1998). DNA was extracted as described above from WT *P. patens*.

The PpRossmann fold 5’ fragment was amplified with primers C265’f and C265’r and the 3’ fragment was amplified with primers C263’f and C263’r (Table 1). Primers for the 5’ fragment were designed to contain 5’ *Nco*I and 3’ *Eco*RI restriction endonuclease sites and the primers for the 3’ fragment were designed to contain 5’ *Eco*RI and 3’ *Not*I restriction endonuclease sites (Table 1). PCR for the 5’ and 3’ fragments was carried out using 0.2 µl (1 U) Phusion polymerase according to the manufacturer’s instructions (ThermoFisher Scientific, Vilnius, Lithuania). PCR products were purified with E.Z.N.A. Gel Extraction kit.

**Table 1.**
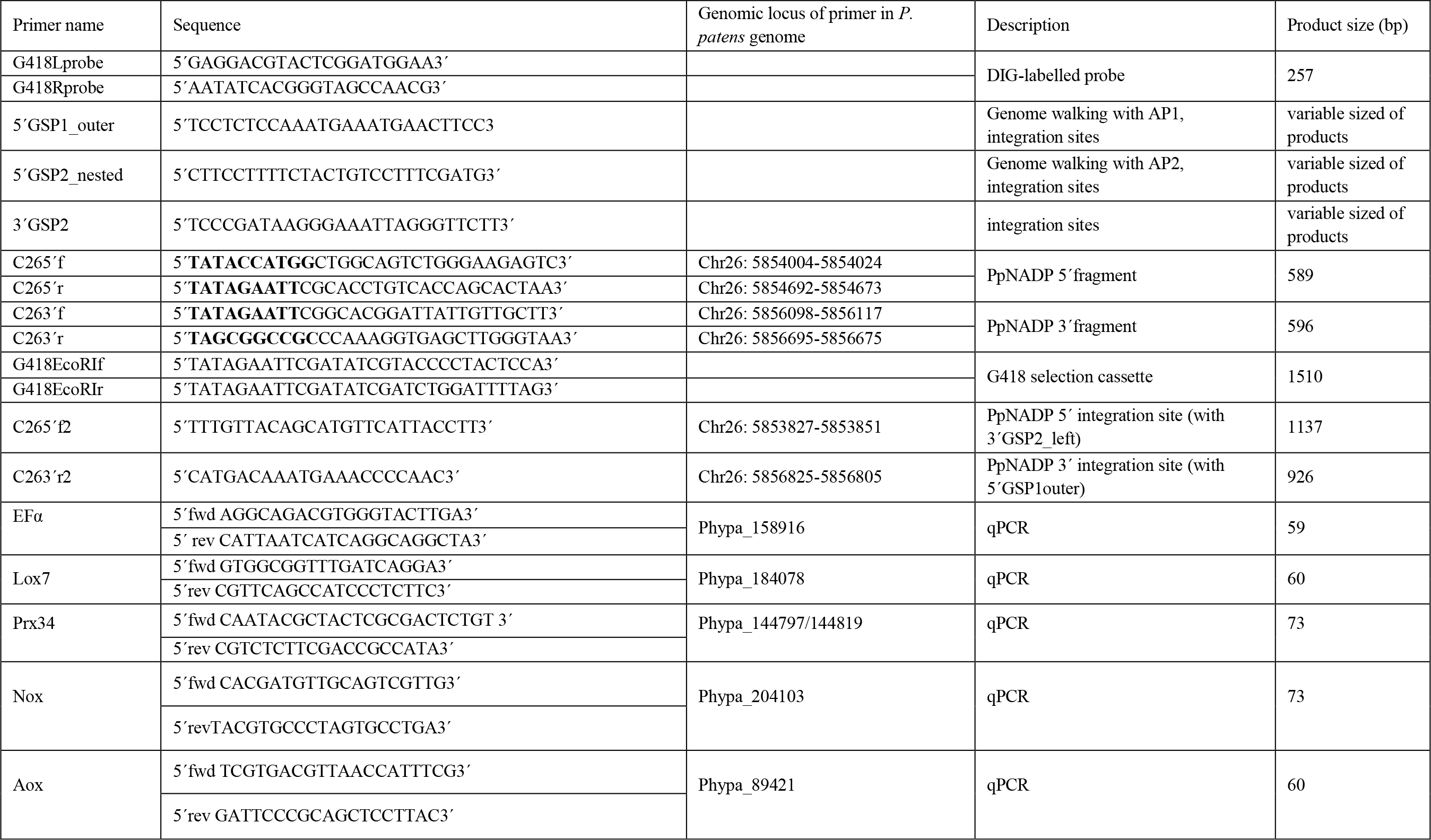
Primers used in the study

The 5’ fragments and pGEM-T easy vector were separately digested with 0.5 µl (10 U) of restriction enzymes *Nco*I and *Eco*RI. Ligation of the digested pGEM-T easy vector and 5’ fragments was carried out overnight at 4 °C using 0.5 µl (1.5 U) of T4 DNA Ligase (Promega). pGEM-T easy vector containing the 5’ fragment (5’-pGEM-T) was transformed into *E*. *coli* DH5α. The 5’-pGEM-T construct was then isolated with GenElute^TM^ Plasmid miniprep kit (Sigma) and digested with 0.5 µl (10 U) *Eco*RI and *Not*I (New England BioLabs). The same restriction enzymes were used to digest the 3’ fragment and to ligate the 3’ fragment and 5’-pGEM-T easy plasmid and the resulting vector containing PpRossmann fold 5’−3’-pGEM-T was transformed into *E*. *coli* DH5α as above.

A plasmid carrying the geneticin G418 resistance gene was obtained from the John Innes Centre, Norwich, UK (http://www.pgreen.ac.uk/). Geneticin selection cassettes were amplified with G418EcoRIf and G418EcoRIr primers having *Eco*RI overhangs (Table 1). Amplified products and 5’−3’ pGEM-T easy vectors were digested with *Eco*RI and ligated to form the knockout constructs. Plasmids were verified by sequencing from the 5’ and 3’ ends. Plasmids were then isolated with Plasmid Maxi kit (Qiagen, Hilden, Germany) and 100 µg of plasmid was digested with 5 µl (100 U) of *Nco*I and *Not*I restriction enzymes and purified with phenol-chloroform extraction in preparation for moss transformation.

### Moss transformation and molecular characterization

Protoplasts were prepared for *P. patens* transformation according to Schaefer et al. (1991). In brief, 300 µl of protoplasts (1 × 10^5^ to 10^6^ protoplasts ml^−1^) were transformed with 10−30 µg of DNA. Transformed protoplasts were plated on cellophane covered BCD plates supplemented with 10 mM CaCl_2_, 45 μM Na_2_-EDTA, 5 mM ammonium tartrate, 6.6% mannitol and 0.5% glucose. The growth conditions were as described for plant material. After 6 d of incubation, protoplasts were transferred onto BCD-growth medium containing ammonium tartrate and geneticin G418 antibiotic (50 µg ml^−1^; Promega) followed by two weeks growth period without the antibiotic and second selection period for two weeks with antibiotic again.

The correct integration of the knockout construct was verified in the moss lines surviving the second selection period. Primer pairs for screening the KO mutant lines were designed to recognize the genomic region upstream of the 5’ integration site and downstream of the 3’ integration site and also for the region of the G418 selection cassette (5’GSP1_outer and 5’GSP2_nested) (Table 1, Fig. S2). The PpRossmann fold 5’ integration site for KO-plants was verified with primer pair C265’2 and 3’GSP2left and the 3’integration site was confirmed with primers C263’R2 and 5’GSP_outer (10 µM each primer).

Three PpRossmann fold KO-plants (lines 17, 24 and 35) that had correct integration sites were chosen for further studies. Ploidy levels of the moss lines were determined based on their nuclear DNA content as measured by flow cytometry at Plant Cytometry Services (Didam, the Netherlands), according to Schween et al. (2003), modified after Heck et al. (2021).

### RNA extraction, cDNA synthesis and quantitative PCR (qPCR)

Two replicates each of the chitosan-treated and untreated PpRossmann fold KO plants and WT were grown in each experiment and used for RNA extraction. Plants were grown as described for the peroxidase activity assay. Chitosan- and water-treated plants were collected 40 min after treatment. Moss tissues were ground in liquid nitrogen and total RNA was extracted according to Chang et al. (1993). Ground moss tissue was transferred to pre-warmed isolation buffer (2% CTAB, 2% PVP-K-30, 100 mM Tris-HCl [pH 8.0], 25 mM EDTA, 2 M NaCl, 0.02% β mercaptoethanol). Samples were extracted three times with chloroform/isoamyl alcohol (24:1), shaken at 250 rpm for 15 min and centrifuged at 10,000 × *g* for 15 min. Samples were precipitated with 8 M LiCl at 4 °C overnight. After centrifugation at 10,000 × *g* for 30 min at 4 °C, the pellet was dissolved in pre-warmed SSTE buffer (1 M NaCl, 0.5% SDS, 10 mM Tris-HCl [pH 8.0], 1 mM EDTA) and extracted once with chloroform/isoamyl alcohol (24:1) as above. The RNA was precipitated from the aqueous phase with two volumes of isopropanol and dissolved in nuclease-free water. The RNA concentration was measured with a spectrophotometer (Nanodrop 2000, Thermo Scientific).

RNA samples from two biological replicates were pooled and total RNA was treated with DNase (1 U/µl, Promega). For cDNA synthesis, the RNA (1 µg) was then incubated with 1 µl (200 ng/µl) random hexamers (ThermoFisher Scientific) at 65 °C for 5 min. Reverse transcription was performed using 4 µl 5× reaction buffer (250 mM Tris-HCl [pH 8.3 at 25 °C], 250 mM KCl, 20 mM MgCl_2_, 50 mM DTT [ThermoFisher Scientific]), 0.5 µl ribolock RNase inhibitor (40 U/µl, ThermoFisher Scientific), 2 µl dNTPs (10 mM each, ThermoFisher Scientific) and 1 µl RevertAid M-MuLV reverse transcriptase (200 U, ThermoFisher Scientific). Reaction mixtures were incubated at 25 °C for 10 min followed by incubation at 42 °C for 1 h. The reaction was stopped by heating at 70 °C for 10 min and the resulting cDNA was diluted with 80 µl of nuclease-free water.

Lipoxygenase7 (LOX7), Peroxidase (Prx34), NADPH-oxidase (Nox) and alternative oxidase (Aox) expression were measured with quantitative PCR (qPCR) according to Lehtonen et al. (2012). Elongation factor EF1α was used as a reference gene. qPCR was carried out using a reaction volume of 10 µl, which included 0.5 µl of forward and reverse primers (5 µM each), 4 µl cDNA and 5 µl of LightCycler 480 SYBR Green I Master mix (Roche Diagnostics GmbH, Germany). Three technical replicates of each sample were included. qPCR was carried out according to the manufacturer’s instructions. The relative expression ratio of the target gene was calculated according to Pfaffl (2001). An analysis of variance (ANOVA; IMB SPSS, v. 25.0, SPSS Inc.) was carried out to determine whether there were significant differences in the expression levels between PpRossmann fold KO-lines (17, 24, 35) and WT *P. patens*.

## Acknowledgements

Doctoral Program in Integrative Life Science (former Viikki Graduate School in Molecular Biosciences), Finnish Cultural Foundation, Future Found of The Faculty of Agriculture and Forestry and Finnish Concordia Found are gratefully acknowledged for financial support. This study was supported by the German Research Foundation (DFG) under Germany’s Excellence Strategy (CIBSS – EXC-2189 – Project ID 390939984).

## Author contribution

Work planning EMM, ELD, RR, JPTV Experimental implementation EMM, PH Manuscript writing EMM, ELD, RR, JPTV

