## Supplemental figures for "Putative NAD(P)-binding Rossmann fold protein is involved in chitosan-induced peroxidase activity and lipoxygenase expression in Physcomitrella"

**Supplementary information**

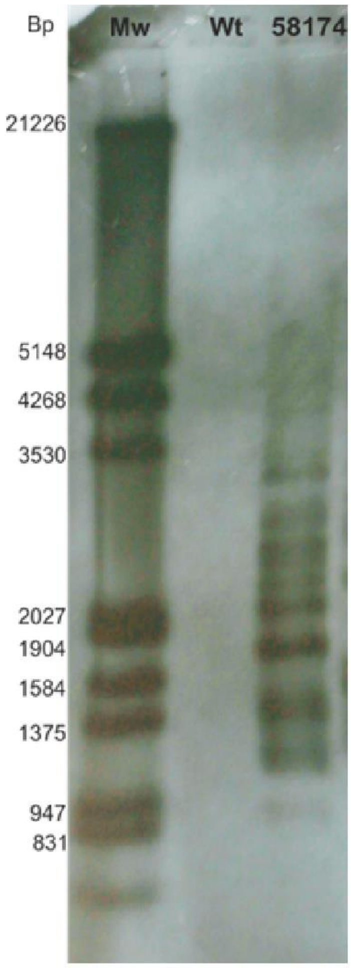


Figure S1. Southern blot analysis for Physcomitrella wild-type and transgenic *P. patens* line 58174. Southern blot analysis with labeled probe complementary to *npt*II indicated many *npt*II cassette integrations within transgenic moss line 58174. A DNA ladder (M) is shown in the first lane of the gel.

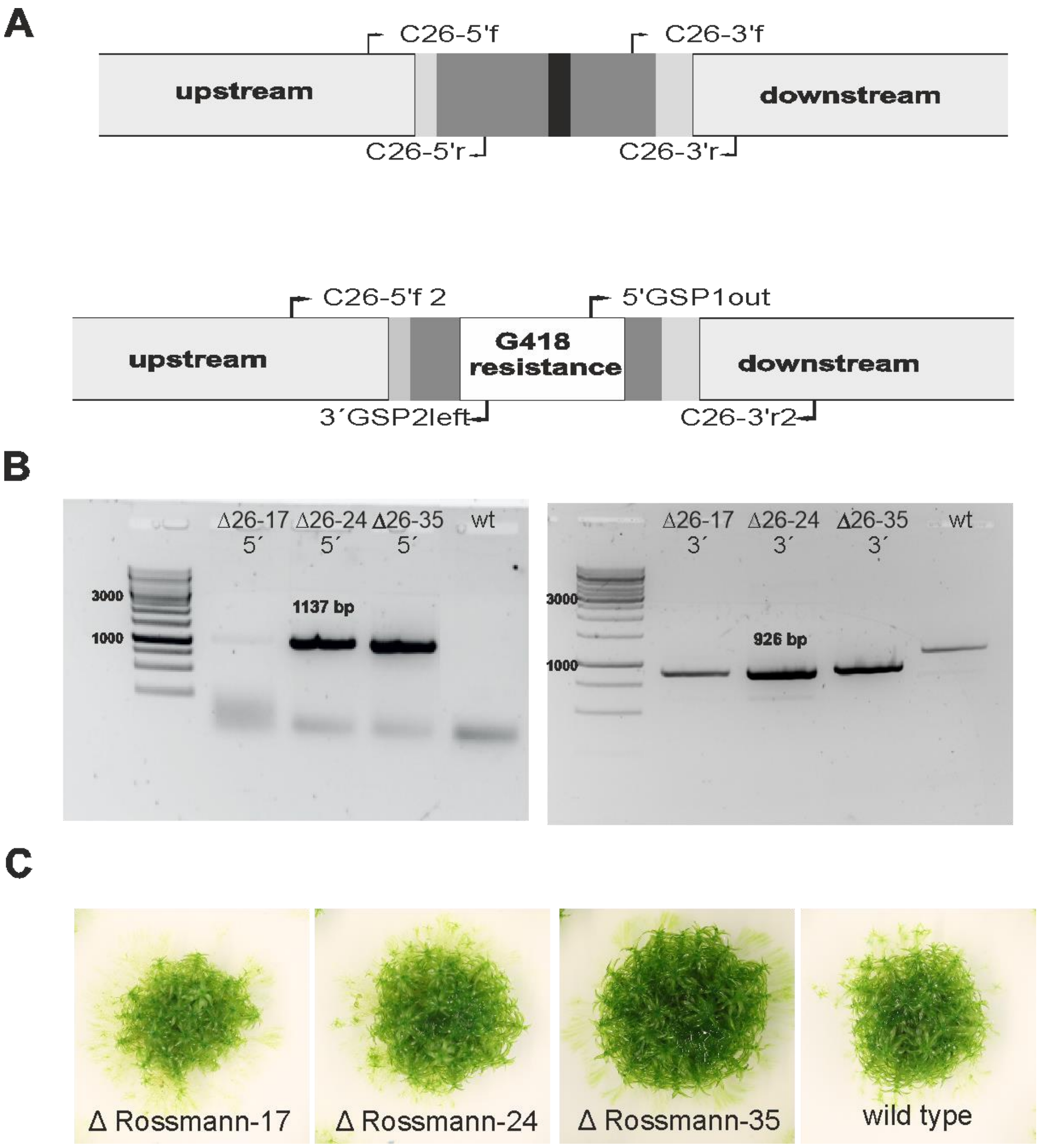


Figure S2. Gene replacement for putative Rossmann fold protein in *P. patens* and the appearance of the resulting transgenic lines on BCD-growth medium supplemented with ammonium tartrate. A) Schematic diagram of the Rossmann fold gene before and after integration of the geneticin G418 resistance cassette. Arrows indicate the sites for primer binding. Dark gray regions represent exons, light gray regions indicate 5´and 3´ untranslated regions and the black region indicates an intron. B) Screening and verification of Rossmann fold KO-lines by PCR indicated the integration of the hygromycin resistance cassette in the place of the gene encoding the putative Rossmann fold protein. C265´f2 and 3´GSP primers were used for 5´integration site and C263´r2 and 5´GSP1outer primers for 3´integration site verification. C) The appearance of knockout and wild-type (WT) plants on BCD-medium supplemented with ammonium tartrate.
